# The coevolution of encephalization and manual dexterity in hominins and other primates

**DOI:** 10.1101/2024.09.17.613411

**Authors:** Joanna Baker, Robert A. Barton, Chris Venditti

## Abstract

The human hand is one of our most remarkable features. We have long, opposable thumbs and a suite of other features argued to be adaptations for interacting with and manipulating our environment, literally extending the reach of our cognitive powers. Consequently, enhanced manipulative dexterity, tool use and increased brain size are considered key features of how our ancestors evolved. This hypothesis predicts that anatomical dexterity and brain size co-evolved, and that this should be evident in the morphology of the hand. To test this hypothesis and to understand how hominin evolution may have deviated from any general primate trend, we collected a dataset of finger length, thumb length, and brain size across 94 extinct and extant species of primates. Using phylogenetic comparative methods, we reveal a strong primate-wide association between brain size and relative thumb length. We further demonstrate that this general association accurately predicts the co-evolution of these traits in hominins. Whilst hominins have significantly long thumbs compared to other primates, we infer that they have arisen from the same underlying evolutionary process acting across the whole primate order: increasing manipulative ability associated with specialized neural control processes. The relationships we recover are consistent with positive feedback between manipulation, tool use and cognition and therefore may go some way towards explaining evolutionary trends towards larger brain sizes among primates, and particularly in hominins. Our results emphasize the role of manipulative abilities in cognitive evolution and emphasize how neural and bodily adaptations are interconnected.

## Main

An enhanced ability for visually-guided grasping and manipulation is often considered one of the defining features of the primate order (1). Primates are also generally considered to have specialized cognitive abilities associated with large brains (2). The relationship between manual dexterity and brain size, however, remains largely unexplored (3, 4).

Within humans, associations between manual dexterity and cognitive function are well established in experimental settings (e.g. 5, 6, 7), and it is often assumed that cognitive ability is linked to tool use and dexterity (3, 8). Tool use is observed in several primate species (9–16), and is in fact but one manifestation of more widespread skills related to extractive foraging (17, 18). Such behaviors should require increased visuo-motor and cognitive skills, such as the capacity to learn and execute complex sequences of actions (3, 19, 20), and have been linked to brain size (14–16, 21). Intriguingly, there is evidence that complex manipulation behaviors have co-evolved with brain size in primates (3, 22), but how might this be reflected in morphological traits that change in response to natural selection and facilitate behaviour? A focus on morphology can bolster our understanding of the importance of the interrelationships of manipulation and neurocognitive evolution. Not only this, but we can also test for them in extinct species like our own ancestors – in which behaviours are unobservable.

An increased ability to manipulate small objects is enhanced by long thumbs (23–25) – particularly relative to the index finger (26). Longer thumbs facilitate diverse grip techniques (27) as well as increasing opposability (28). While humans have refined precision grasping ability (28, 29), there are varying degrees of opposability across primates (26, 28) and precision grasping behaviours are found even in species with only pseudo-opposability, such as capuchins (23, 30). If, as seems likely, fine manipulative abilities require enhanced sensory-motor control, then we would expect to see covariation between thumb length and brain size. Indeed, this might explain some of the marked variation in relative brain size observed across primates and the tendency for it to increase through time (31).

We predict a positive relationship between relative thumb length and brain size across primates. (Figure 1a, solid line). No relationship (Figure 1a, dashed line) might indicate that manipulation had negligible consequences for brain size - or that multiple features combine with thumb length to increase manipulative potential (23, 32).

**Figure 1.**
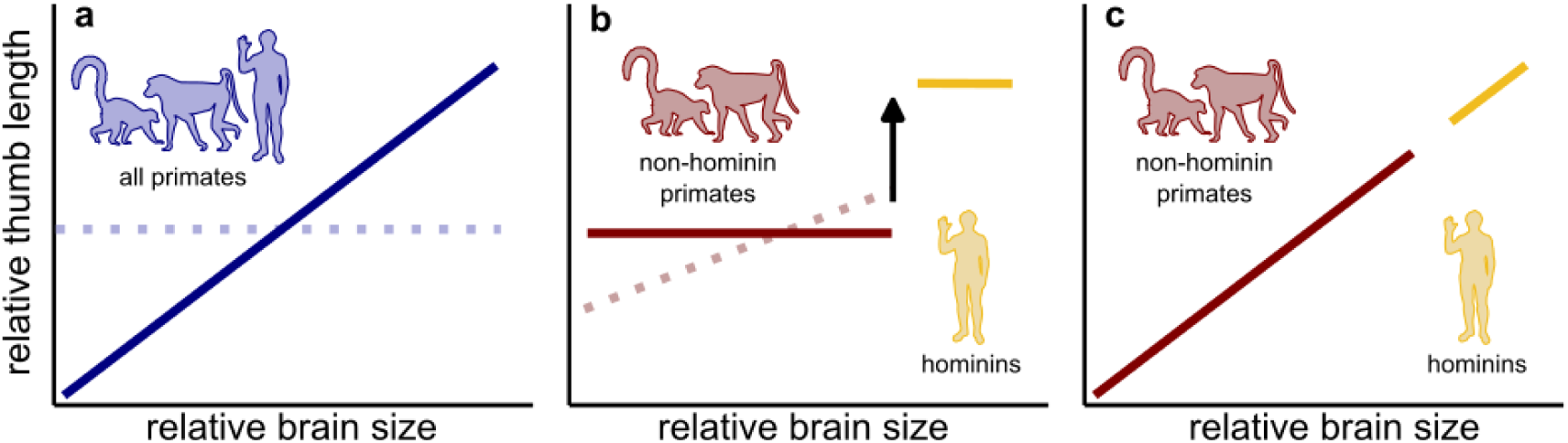
Expectations for the relationship between thumb length and brain size across all primates and how this could be affected by the inclusion of hominins. Across all primates (a), we could observe a significant positive relationship (solid line), or no relationship (dashed line); we do not expect a negative relationship. However, this positive relationship could be being driven or affected by the inclusion of hominins (b). If hominins have a large shift in thumb length, then a spurious positive relationship may be detected across all primates. In this case, we would expect no relationship among other primates (solid red line, b). However, if the relationship is maintained – or a different relationship altogether exists (dashed red line, b) alongside a shift in thumb length in some or all hominins, it implies some increase in thumb length beyond that explained by manual dexterity and motor control. Finally, it is possible that the long thumbs and large brains of hominins arose as a part of the same process across all primates (c). In our models, we cannot estimate a separate slope for hominins owing to their small sample size (n = 6), hence here they are depicted as a mean shift with no slope (yellow line, b).

Hominins have much greater relative thumb length compared to other apes (30, 33), which has specifically been linked to refined precision grasping (23, 29, 32). The timing of the emergence of such behaviors and associated morphologies, however, are highly contested (24, 34–37). If long thumbs are specifically advantageous for grips facilitating hominin tool use and manufacture, then we would expect an increase in thumb length in species that habitually use and manufacture tools, but no association amongst any other species (Figure 1b, solid lines). If, instead, we identify an overall positive primate association, where some hominins are outliers (e.g. dashed line, Figure 1b) this implies a physical ability for high dexterity without any need for additional neural processing capacity. In such a scenario, we might look to other factors that are likely to have driven selection on hand morphology besides dexterity. Alternatively, both large brains and long thumbs in hominins may have arisen as an extreme manifestation of the same general evolutionary process in primates. In this scenario, we would observe a relationship between thumb length and brain size across all primates which also quantitatively predicts the co-evolution of these traits in hominins (Figure 1c). This would be in line with suggestions that brain size variation in primates reflects sensory-motor control (19) and many morphological adaptations associated with dexterity pre-date hominins (30).

Here, we test whether brain size and manual dexterity (indexed by intrinsic relative thumb length, i.e. accounting for finger length) were linked during primate evolution using comparative phylogenetic analysis and a dataset of 94 fossil and contemporary primate species (Figure 2). We examine whether hominins have exceptionally long thumbs in the context of statistical phylogenetic outlier tests. Finally, we assess whether any hominin outliers in terms of thumb length are explained by selection for improved cognition associated with manipulative ability.

**Figure 2.**
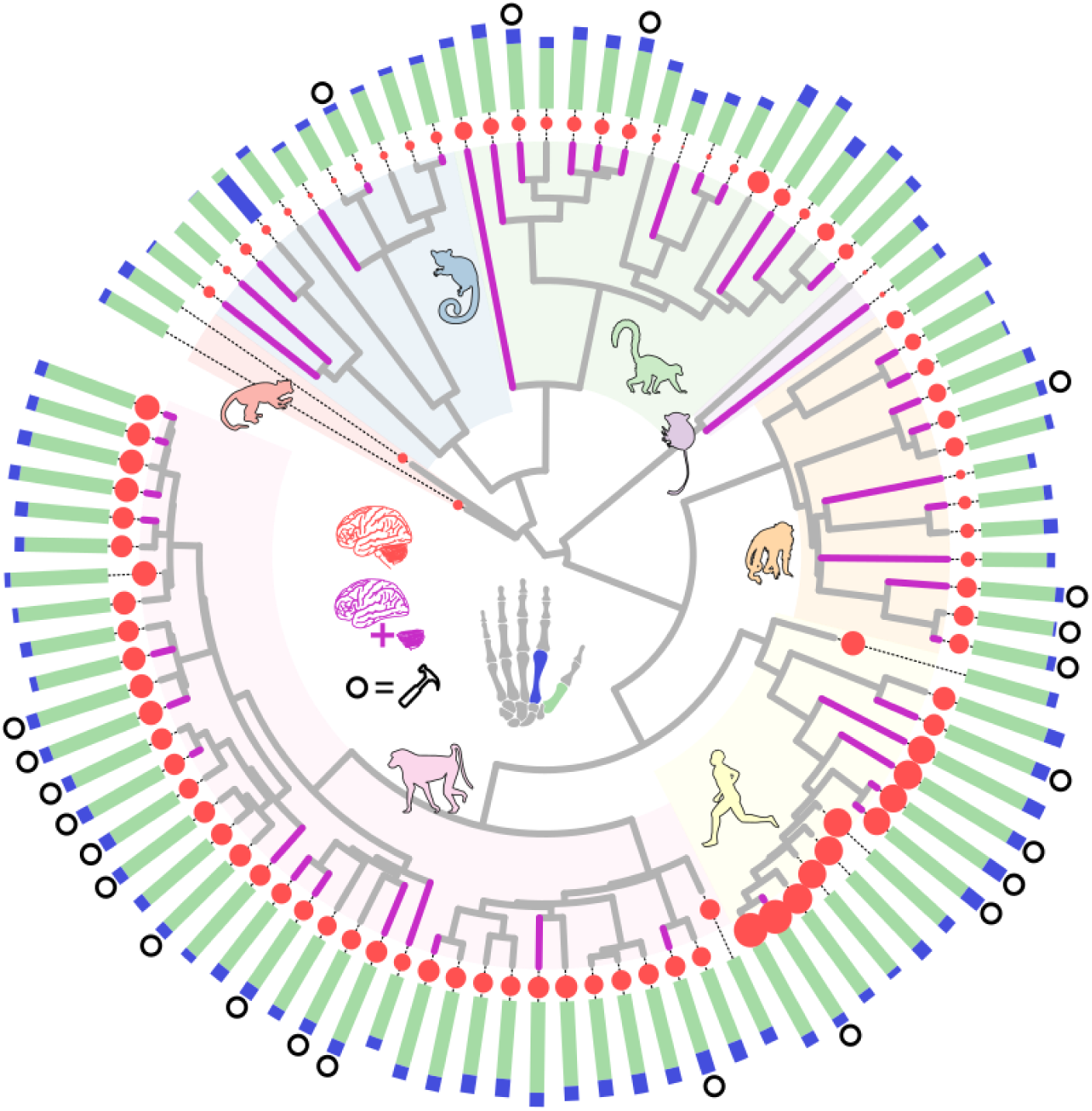
Data used to test the relationship between dexterity and cognition. Manual dexterity is measured using the relationship between the length of the first metacarpal (green) and the second metacarpal (blue) – the length of each of these bones is shown by the bars at the tips of the tree (shorter bone superimposed on top of the other). Whole brain size is represented by red circles at the tips of the tree. Species for which we have both cerebellum and neocortex volumes are indicated in purple. Species with any documented tool-use are denoted by open circles at the tips. Major clades of the primate radiation are highlighted. Silhouettes are not to scale.

### The uniquely long thumbs of hominins

Although features of the human hand, such as long thumbs, pre-date the origin of systematic tool production (30, 34, 35), features such as long thumbs are repeatedly highlighted as indicative of increased manual dexterity (23, 29, 32) and are even sometimes used as an marker for tool culture (8, 32) – (reviewed in 38). We set out to assess whether hominin thumbs are truly unique relative to all other primate species – that is, are humans unique in terms of their intrinsic hand proportions (30)?

Using phylogenetic generalized least squares (PGLS) regression models implemented in BayesTraits (39) and accounting for phylogenetic uncertainty by using a sample of dated trees (see Methods), we find that thumb length and finger length are strongly linked across all primates (n = 94, *finger-only* model, Figure 3a). The relationship is significant in 100% of our tree sample (see Methods), with a median slope parameter (ϐ_[finger]_) = 0.88. There is also high phylogenetic signal (median λ=0.86). The median R^2^ of the model is 0.88. When we remove hominins from the analysis, the relationship remains strongly significant (n = 88, median ϐ_[finger]_ 0.90, p_x_ < 0.05 in 100% of our sample) with high phylogenetic signal (median λ = 0.88).

**Figure 3.**
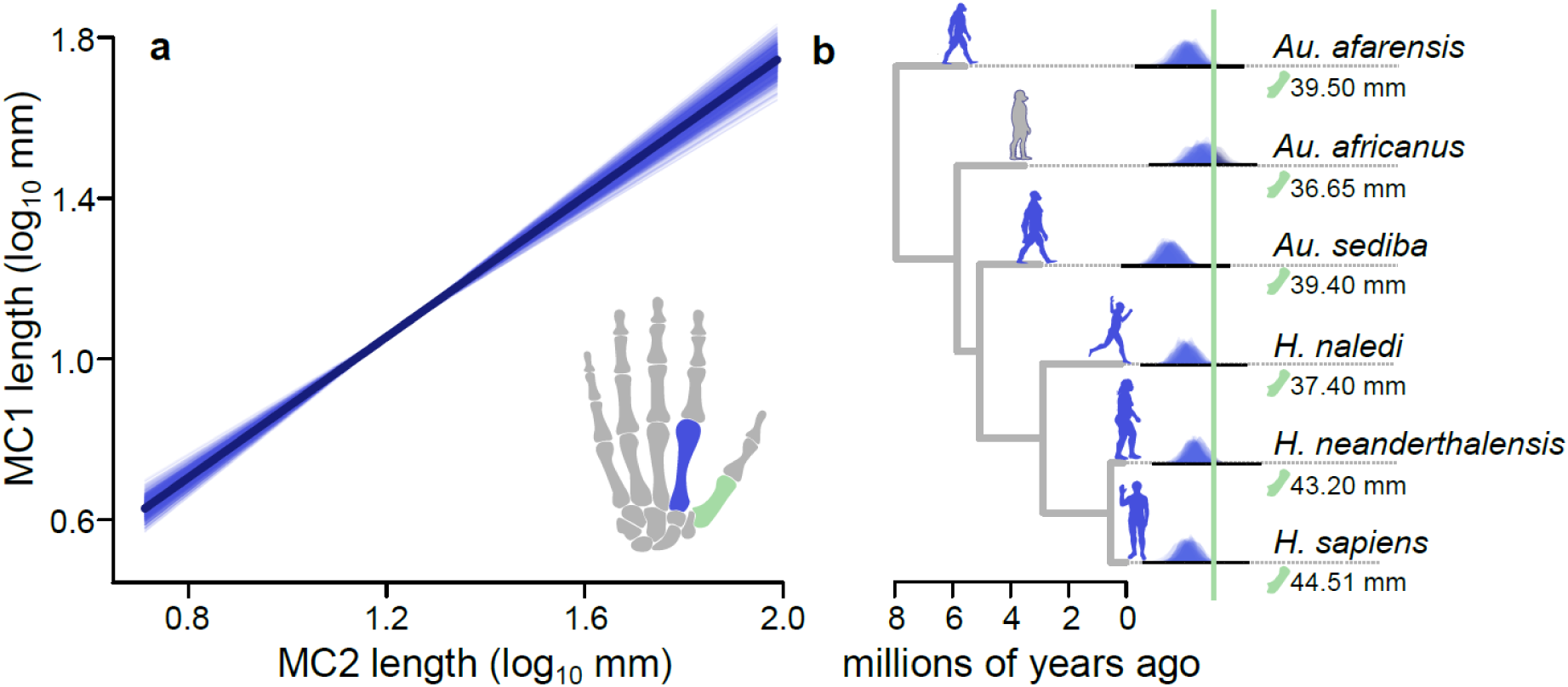
The relationship between finger length and thumb length across primates. (a) The predicted slopes for the inferred relationship for our finger-only model are plotted for 25 random samples from fifty random trees across the sample. The median fitted relationship is superimposed in dark blue. (b) The hominin phylogeny (using a single representative from the sample) is plotted along with the posterior distributions of imputed thumb lengths from the finger-only (blue) model. There are 100 distributions for each hominin for each model – one for each of the topologies in the sample. The real thumb length of each species is indicated by the green line. Silhouettes are shown for representative purposes only and are not to scale.

Given the finger-only model estimated across all non-hominin primates, we find that all but one hominin species (*Australopithecus africanus*) is a significant outlier (Figure 3b, identified using phylogenetic outlier tests – see methods). In general, the relationship across all primates predicts hominin species to have thumb lengths that are significantly much too small: as expected, hominin thumbs are significantly longer than those of other primates (Figure 3b), supporting the hypothesis that hominins underwent adaptive evolution for enhanced dexterity.

### The co-evolution of brain size and thumb length

We find a significant positive relationship between brain size and thumb length, accounting for allometric effects (finger length): *whole-brain* model, median β_[brain]_ = 0.120, p_x_ < 0.05 in 100% of trees, Figure 4, Table 1. In this whole-brain model, the relationship between thumb length and finger length is maintained (median β_[finger]_ = 0.701, p_x_ < 0.05 in 100% of trees), and there is high phylogenetic signal (median λ = 0.830). We then repeated our whole-brain model excluding all hominins (n = 6) to determine whether their inclusion was affecting the observations across all primates (e.g. Figure 1b). We find that there is still a significantly positive relationship between relative thumb length and brain size (median β_[brain]_ = 0.094, p_x_ < 0.05 in 100% of trees) as well as thumb length and finger length (median β_[finger]_ = 0.760, p_x_ < 0.05 in 100% of trees) across all non-hominin primates. That is, hominins are not driving the observed association between thumb length and brain size (Figure 1).

**Figure 4.**
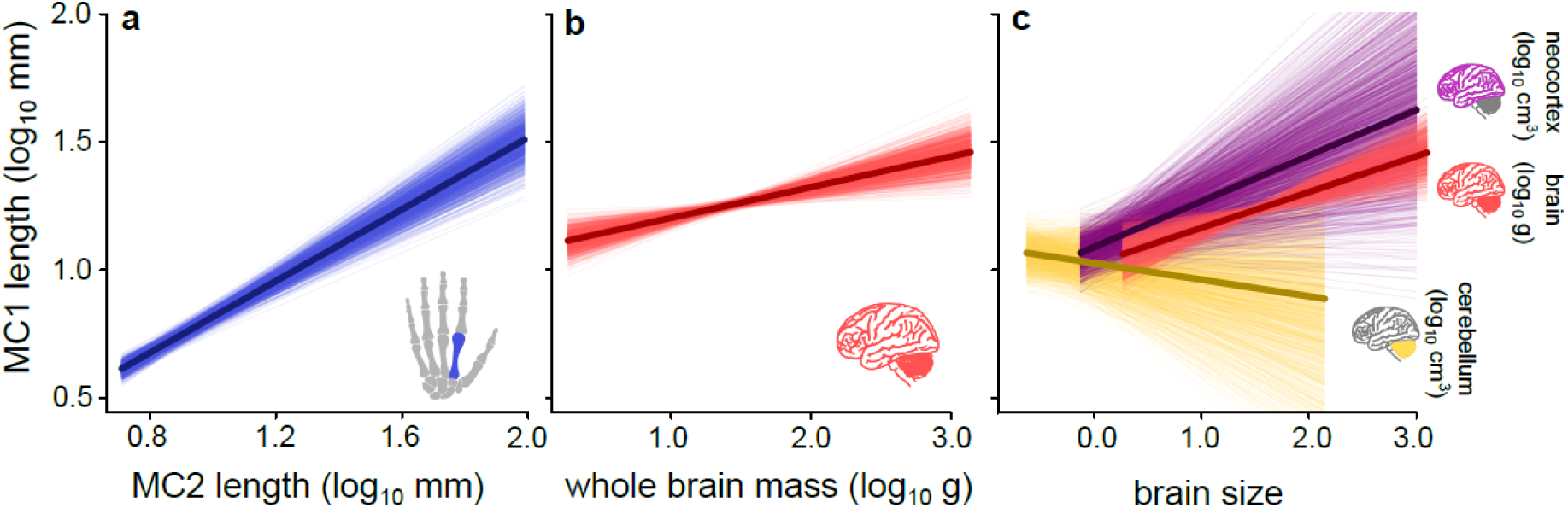
The relationship between thumb length, finger length and brain size across primates from our whole-brain and brain-regions models. In all panels, we depict a random (N = 25) sample of predicted relationships from a random 50 trees in our sample, holding the unplotted variable at its mean value. The median predicted relationship is calculated across all trees and is shown by a darker line. (a) The predicted relationship between thumb and finger length estimated across our full sample (N = 95) in our whole-brain model (mc1 ∼ mc2 + brain). (b) The predicted relationship between thumb and whole brain mass in our whole-brain model. (c) The relationships estimated from our brain-regions model (mc1 ∼ mc2 + neocortex + cerebellum) using N = 49 extant primates. All plotted slopes are significant except for the relationship between cerebellum volume and thumb length (yellow lines, panel c).

**Table 1.** Regression parameters for all models with and without hominins (the range of medians per tree in the sample). The percentage of the sample in which we observe significance (p_x_ < 0.05) is reported in square brackets after each parameter.

| Model | N | median $R^2$ | median $\beta_{\text{finger}}$ | median $\beta_{\text{brain}}$ | median $\lambda$ |
| --- | --- | --- | --- | --- | --- |
| finger-only | 94 | 0.87-0.89 | 0.87-0.89 | n/a | 0.82-0.89 |
| finger-only (excl. hominins) | 88 | 0.88-0.89 | 0.89-0.91 | n/a | 0.84-0.91 |
| whole-brain | 94 | 0.85-0.86 | 0.69-0.72 [100%] | 0.11-0.13 [100%] | 0.78-0.87 |
| whole-brain (excl. hominins) | 88 | 0.86-0.88 | 0.74-0.78 [100%] | 0.08-0.11 [100%] | 0.84-0.91 |
| brain-regions | 49 | 0.91-0.92 | 0.72-0.76 [100%] | neo: 0.16-0.20 [100%]<br>cereb: -0.08 - (-0.04) [0%] | 0.92-0.97 |
| brain-regions (excl. human) | 48 | 0.91-0.93 | 0.79-0.82 [100%] | neo: 0.13-0.17 [96%]<br>cereb: -0.10 – (-0.05) [0%] | 0.92-0.97 |

Improvements in fine-grained visuo-motor processes such as visually guided manipulation are expected to be associated with expansion of brain regions mediating these processes. Substantial areas of the primate neocortex and cerebellum are involved in visuo-motor control, and coordinated expansion of these structures explains much of the variation in brain size among primates (19). In a reduced sample of primate species with data on neocortex and cerebellum volume (n = 49, Figure 2), we identified a significant positive relationship between thumb length and both finger length (β_[finger]_ = 0.735, p_x_ < 0.05 in 100% of trees) and neocortex (Figure 5, β_[neocortex]_ = 0.178, p_x_ < 0.05 in 100% of trees, see supporting information and Table S2 for full results). We refer to this model as our *brain-regions* model – which tests the effects of both brain regions simultaneously. Our results are qualitatively identical without humans (Table 1), indicating that such observations may not be limited to our own species.

**Figure 5.**
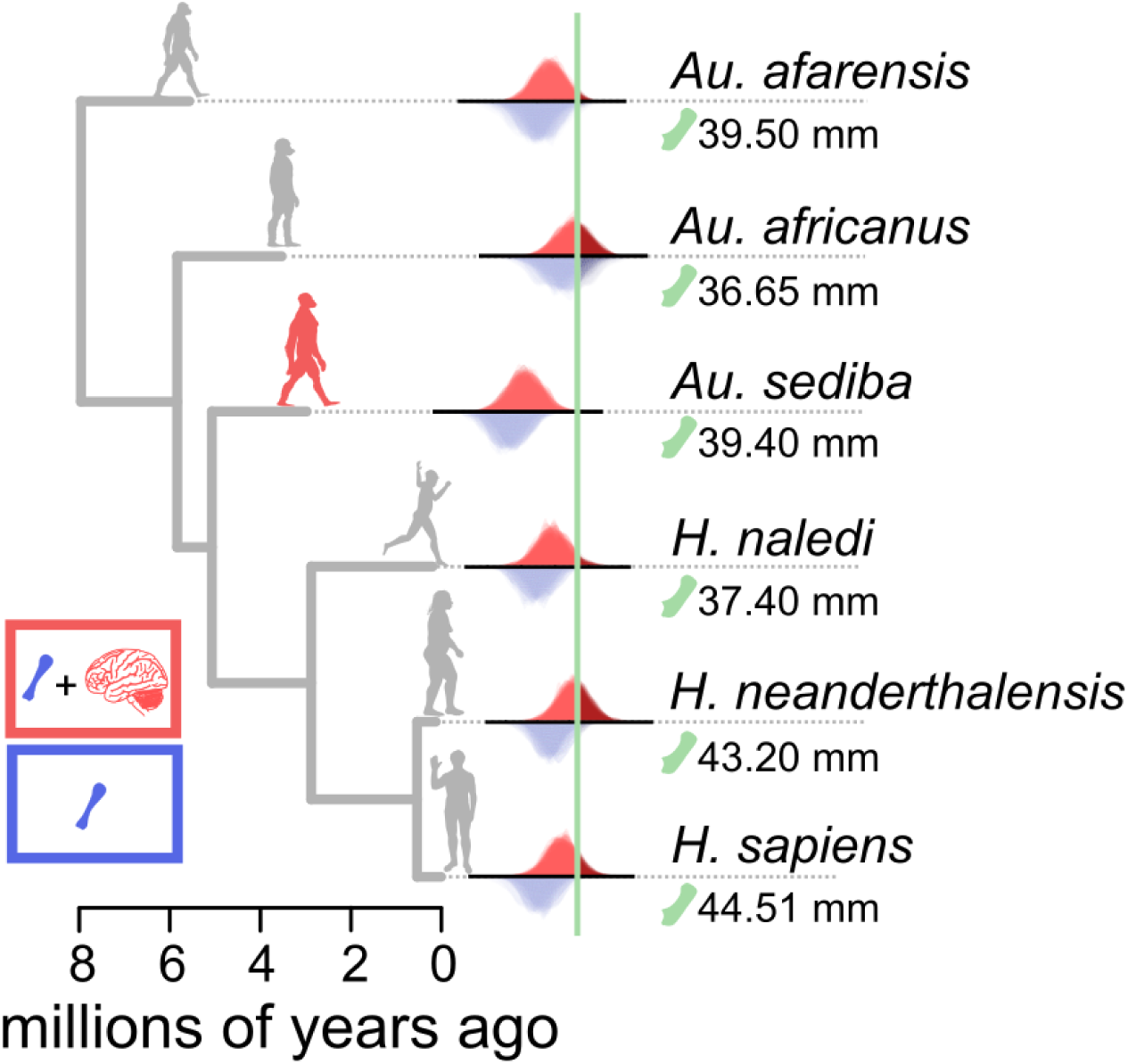
No difference in thumb length amongst hominins after accounting for brain size. Hominins are not outliers to the relationship between thumb length and brain size (except for Australopithecus sediba). Here we show the hominin phylogeny (using a single representative from the sample) and the posterior distributions of imputed thumb lengths from the whole-brain (red) and finger-only (blue) models. There are one hundred distributions for each hominin for each model – one for each topology in the sample. The real thumb length of each species is indicated by the green line. Silhouettes are representative only and not to scale.

There is no significant association between thumb length and cerebellum volume (Figure 5) – nor do we find any link with an anatomical measure of binocular vision, convergence of the orbits (supplementary material). This is surprising, given the established role of the cerebellum and cortico-cerebellar networks in fine visuo-motor control and management of complex behavioral sequences (19, 40). Our brain-region relationships are more variable than those observed for the whole-brain (see Figure 5) and are also potentially affected by small sample sizes. However, the conclusions are not affected by the exclusion of the apes, which demonstrate rapid cerebellum expansion (41) nor do they diverge between haplorrhines and strepsirrhines despite relative differences in the sizes of their relative brain regions (42). We therefore reveal the intriguing possibility that neocortex, but not cerebellum, is implicated in thumb length evolution across primates. Although we did not predict this dissociation, a cortical contribution is in line with experimental evidence from modern humans, suggesting that motor cortex functioning and grey matter volume are both linked with manual dexterity and hand control (43–45) Further attention is required to untangle which cortical regions are associated with thumb length and what neural processes were involved.

### Hominin thumbs, refined precision grasping, and tool use

Some authors suggest that hominins diverged from other primates in evolving a uniquely specialized morphology adapted for the manufacture and habitual use of tools (35, 36). However, we find a primate-wide association between brain size and thumb length, indicating that thumb length is a more general measure of dexterity not specific to humans. To test whether the association across primates also predicts co-evolution of these traits in hominins, we conducted another set of phylogenetic outlier tests (e.g. 46) this time using our whole-brain and brain-region models estimated across all non-hominin primates.

After accounting for brain size, no hominin species is identified as an outlier to the thumb length and brain size relationship across all other non-hominin primates (Figure 5, red distributions). This is to the exception of *Australopithecus sediba,* in which thumb length is much greater than would be predicted by its brain size. In all but *A. sediba*, the posterior distribution of estimated thumb lengths overlaps the true value by more than 5%, in more than 95% of trees. Overall, hominins conform to the expectations given by the relationship between thumb length and brain size for all other primates (see Figure 1c). This is striking: whilst hominin thumbs are outliers amongst primates in terms of length (Figure 3b), this is almost entirely explained by their brain size (Figure 5).

As with our results for intrinsic hand proportions, we find a general relationship across primates between relative thumb length and brain size: Most hominins have dexterous abilities predicted by their cranial capacity. For example, although *Australopithecus africanus* does not have significantly long thumbs relative to finger length (Figure 4), it still conforms to the whole-brain relationship observed across all primates. That is, the combination of brain size and thumb lengths in this taxon leads to manipulative ability comparable to other hominins – as suggested by other studies (26, 35). Whilst hominins do have long thumbs, this arises from a common underlying process linking thumb length to brain size across the primate order: selection on manipulative ability.

The hypothesis that longer thumbs are specifically advantageous with regards to tool use predicts difference in thumb length between tool-using primates and those which have not been observed to use tools. To test this prediction, we used a comprehensive compilation of observed tool-use (47) in combination with PGLS models to assess the relationship between thumb length and tool use, accounting for both finger length and phylogeny (see methods and supporting information). In this model, we find that there is no significant difference in thumb length between tool-users and non tool-users (p_x_ > 0.05) in any of the 100 trees. There has been some considerable debate surrounding the definition of tool-use e.g.(47–49). However, our results are consistent when we use alternative definitions of tool use: firstly, in species observed to use ‘true’ tools – where objects are explicitly manipulated out of their original context (48, 50); and secondly, in species observed to explicitly manufacture or modify objects prior to use (47, 49). We find the same result even after removing species where either only a single individual has been observed using or making tools, or where observations came only from captive animals (47). Accounting for brain size does not change this result.

In short, there is no difference in thumb length among species that use tools – at least not among extant primates. The debate on which extinct apes and hominins are likely to have been capable of human-like precision grasping behaviours is extensive and ongoing (26, 29, 35, 51, 52). Our results highlight that, in isolation, no single morphological feature – including thumb length – is likely to be informative as to the tool-making behaviour of individual species (32). Instead, it is more likely that a suite of complex characteristics and morphologies (32, 38) – including the increased capacity to understand complex behavioural sequences and their consequences – has led to the remarkable dexterity observed in our own species.

### The curious case of Australopithecus sediba

The only hominin who does not conform to the whole-brain relationship observed across all primates is *Australopithecus sediba.* The thumb length of this species remains an outlier amongst primates even after accounting for brain size (Figure 4) and thus conforms to the unexpected scenario outlined in Figure 1b. Our results demonstrate that *Australopithecus sediba* has substantially longer thumbs than predicted by its brain size (Figure 4) – indeed, MH2, the skeletal specimen representing *Australopithecus sediba,* is noted to have an unusually long thumb (26). At face value, this would imply that this species has greater dexterity than other hominins without the associated neural investment. However, the estimated potential manipulative ability of *Australopithecus sediba* falls within and does not exceed the range of modern humans (26). While it could be that our estimates of thumb length or brain size are incorrect for this species, overall, the morphology of MH2 suggests hand use distinct from other hominins (53) – including earlier species*. Australopithecus sediba* possessed a repertoire of adaptations linked to both its own form of arboreal locomotion along with dexterous manipulation (33, 53–55) – likely underpinning the cause of this species’ deviation from the general primate relationship.

### Concluding remarks

It has been argued that (56): “*To reach and grasp an object with a hand is a huge achievement… the refinement in sensorimotor control within reaching space that such dexterity requires is central to understanding human cognition in general*”. Our results corroborate this view, and the broader point that appreciating the links between bodily and brain adaptations is key to understanding neuro-cognitive evolution. The significant positive association between relative thumb length and brain size across primates highlights an important role of the ability to manipulate food, objects, or the environment in brain size evolution, and emphasizes the neocortical contribution to these behaviours. The generality of the relationship between relative thumb length and brain size can go some way to explaining the so-called uniquely hominin condition of having long thumbs. A more complete picture might emerge with the increasing availability of more data for other, particularly earlier, hominins. Further to the relationships we reveal here, uncovering links between cognition and additional morphological features associated with dexterity e.g. (32, 38) may allow us to untangle the nuanced picture regarding the suites of traits associated with hominin tool use (26, 32) and their origins.

## Materials and methods

### Primate phylogeny

All of our analyses are performed on a random sample of 100 of the most-parsimonious topologies obtained from the recently published comprehensive Euarchonta phylogeny including 894 fossil and extant primates (57). As the original sample of trees is not time-calibrated, we dated these topologies using a tip-dating procedure adapted from the original paper (57) and implemented in BEAST v2.7 (58). For full details on our tip-dating procedure, see the supplementary material. As BEAST is implemented in a Bayesian framework, it gives a posterior distribution of dated trees for each of the 100 topologies in our sample. We created a single representative phylogeny for each topology by calculating a median tree based on the Kendall-Colijn distance metric (59). We did this using the treespace library (60) in R v.4.0 (61). All our downstream analyses are performed on the sample of 100 median dated trees which are provided as supplementary data to this paper.

### Morphological data

We collected data on metacarpal measurements (lengths in millimeters) for primate species from the literature. We only included species found in the phylogeny (57). We preferred compilation estimates (i.e., species-level data) but where specimen level data were included, we took a weighted average across all specimens (weighted, where possible, by number of specimens measured). Where individual specimens were measured by multiple sources, we preferred, arbitrarily, the most recently published source for each specimen. Our final dataset included finger bone measurements spanning 168 primate species, including 8 hominins. A full list of measurements and their sources can be found in the supporting information. Here, we use the first and second metacarpals as proxies for thumb and finger length respectively to measure relative thumb length and intrinsic hand proportions. However, our results remain qualitatively identical when phalanges are used instead – or if we use the 3^rd^ or 4^th^ digit instead of the 2^nd^ digit (supplementary material). Additionally, the metacarpals are a robust and reliable indicator of overall digit length in our sample (supplementary material). We prefer to use bone length over recently proposed kinematic models (26) for measuring manipulative ability as these are directly measurable quantities which are likely to face direct selection pressure from the environment, although our conclusions remain robust even when using these metrics (see supplementary material).

We then collected brain mass data for these species. Brain masses or volumes were taken from the literature (see supporting information for full list of sources). In some cases – mostly for fossil taxa – we converted endocranial volumes to masses. Whilst endocranial volume is often converted to brain mass using the specific gravity of brain mass (1.036 g/mL)(62–64), the majority of our extant data sample comes from a paper which uses a conversion of 1 g to 1 cm^3^ – and does not record which values were volume conversions (65). For consistency, therefore, we use this conversion where necessary. The species for which this was done are recorded in the supporting information.

Tool-use, true tool-use and tool manufacture are taken from a published and comprehensive compilation (47). Any species not included in this compilation were assumed to have not been observed using tools. This data is limited to only extant taxa. Our final dataset is graphically represented in Figure 2.

### Phylogenetic comparative analysis

We use phylogenetic generalized least squares (PGLS) multivariate regression models implemented within a Bayesian Markov Chain Monte-Carlo (MCMC) framework to test for an association between manual dexterity and cognitive ability. In all models, our response variable is the length of the first metacarpal and the second metacarpal length is included as a predictor to measure intrinsic relative thumb length. For our whole brain models, we included brain mass as an additional covariate, and for our brain region models we included neocortex volume and cerebellum volume. We additionally test each of the brain regions separately to ensure that our results are not skewed by collinearity between predictors.

For all models, results are summarized across the sample of trees where the model is run separately for each tree. We assess significance of the parameters using two criteria: Firstly, the proportion of the posterior distribution that crosses zero (p_x_); where this proportion ≤ 0.05, we consider a variable to be significantly different from zero.

Secondly, the first criterion must be met in at least 95% of topologies for us to consider a variable as significant. For comparison, we summarize parameter estimates using median values – and to summarize across all trees, we then take the median of those, which we refer to as β_[x]med._

All the variables we include in our models are significantly associated with body size and including body size as a covariate may introduce issues with multicollinearity.

Additionally, when included into our models, body size was non-significant in 98% of topologies. Our results remain qualitatively identical when body size is included, and so here we present our results without body size.

### Phylogenetic imputation and outlier tests

To identify which hominins (if any) were outliers in terms of their intrinsic hand proportions, we conducted a phylogenetic outlier test (e.g. 46) using a phylogenetic imputation procedure (39) to predict hominin thumb lengths. This predictive modelling approach simultaneously incorporates the parameters of a regression model as well as the phylogenetic position of each taxon we wish to predict for. As with all our main analyses, the imputations are calculated using PGLS regression models implemented within a Bayesian MCMC framework. We estimate the thumb length for each hominin given the parameters of each of our three regression models (finger-only, whole-brain, and brain-regions) calculated across the rest of primates. We then assess whether each species is an outlier using the full distribution of predicted values for thumb length– where the distribution overlaps the true value by less than 5% in more than 95% of trees, it can be considered a phylogenetic outlier.

## Acknowledgments

This work was supported by Leverhulme Research Leadership Award RL-2019-012 to CV. We would like to thank Thomas Püschel and Suzy White for helpful insights and discussion regarding the paper. We are grateful to both Campbell Rolian and Pierre Lemelin for providing their data for us to use in our analyses. Credit and thanks to James Bowden for drawing the Adapiformes silhouette (Figure 1).

## Notes

### Competing Interest Statement

The authors have declared no competing interest.

