## Supporting Information for "The coevolution of encephalization and manual dexterity in hominins and other primates"

### Dataset Collection

Data was collected from the literature on the length of the proximal phalanges, intermediate phalanges, and metacarpals of digits 1-4. We did not include the 5th digit as this is less involved in human precision finger grips (1) and there is generally less data available. We focused our search on the first and second metacarpals.

We collected the lowest possible taxonomic designation for the record as provided by the original source. Where species names were not provided, but we had information on specimens, we looked up the specimen ID in the original collection for additional taxonomic information. For example, ID UM_101963 is recorded as *Carpolestes* in the finger-length source (2), but the specimen record on MorphoSource records the full species name as *Carpolestes simpsoni*, associated with the original publication (3).

We resolved duplicates in the following ways. Firstly, if there were any duplicates identified within an individual source, we took an average. Next, we addressed duplicate measurements across sources – and for these, we chose a representative value rather than taking an average. This is because we cannot always be certain of which specimens were included in the calculation of the original published values. In these cases, we selected the most recently published estimates. Where multiple estimates were published in the most recent year, we selected the first alphabetically. Finally, we resolved duplicates within a species by taking an average of multiple specimens but prioritised using already published taxon-level estimates. This resulted in a dataset with 177 individual unique taxa with length on at least one of the finger bones (Figure S1, Supporting Data S1).

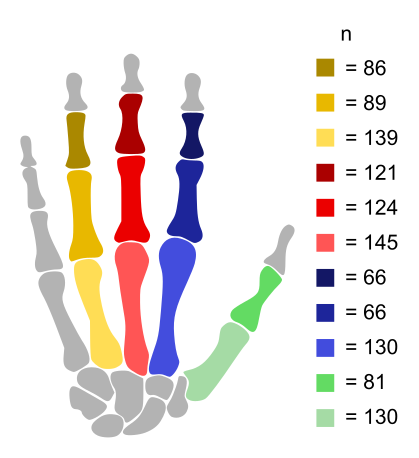
**Figure S1. A schematic representation of the finger bone length dataset used in this study.** Sample sizes are non-overlapping (i.e. not all species have all measurements).

### Tip-Dating Procedure

All of our analyses are performed on a random sample of 100 of the most-parsimonious topologies obtained from the recently published comprehensive Euarchonta phylogeny including 894 fossil and extant primates (4). As the original sample of trees is not time-calibrated, we dated these topologies using a tip-dating procedure adapted from the original paper (4) and implemented in BEAST v2.7 (5).

Before dating, we removed several species owing to uncertain placement and/or taxonomic affiliation (Cercopithecini sp. indet. AUH 1321, Colobinae indet. KNM-BN 1251, Colobinae indet. KNM-TH 48368, Cheracebus purinus, Tupaia sp. UNSM 87244, and Dermoptera indet. Pkg 240 and Pkg 335). Our preliminary analyses demonstrated that inclusion of taxa with broad date ranges and uncertain placement resulted highly variable node ages depending on which clade the taxa fell within. We chose to exclude these species and therefore removed the influence of such taxa on our divergence dating analyses.

We then estimated branch length date variation by fixing the topology. We conditioned the fossilized birth-death process (6) on the root, and placed a uniform root calibration prior of between 66 and 130 million years as described in Dos Reis et al (7). We used an optimized relaxed clock with the mean clock rate restricted to vary between 0.01 and 0.02 (just outside the ranges observed in the maximum credibility tree dated by Wisniewski et al (4) to facilitate faster convergence (though results are identical if this parameter is allowed to freely vary). The σ parameter of the log-normal distribution of the clock rates is drawn from an exponential prior with a mean = 1. Transition and transversion rates were drawn from a gamma prior (α= 0.2, β = 0.5) and (α= 0.2, β = 0.25) respectively. Finally, a uniform prior ranging between 0 and 1 was placed on the turnover rate and for the sampling proportion, we used a beta distribution (α = 5, β=90). All polytomies were randomly resolved before tip-dating but were collapsed afterwards back to the original topology.

We then created a single representative phylogeny for each topology by calculating a median tree based on the Kendall-Colijn distance metric(8). We did this using the treespace library (9) in R v.4.0 (10). We also, for our supplementary analyses (see below) and visualization purposes, created a single overall reference tree by calculating an additional median tree based on our sample of trees.

### Using metacarpal length as a proxy for finger length

Generally speaking, one of the features associated with pad-to-pad precision grasping is a long thumb relative to the lengths of other fingers (11). Specifically, the ratio of thumb to index finger is often used as a proxy for manipulation ability and dexterity (12, 13). To maximize our sample size, we focus on using only a single bone from the thumb and the index finger, the metacarpals. Metacarpals are significant predictors of both adult and foetal body size (14, 15) and have previously been used as a proxy for finger and thumb lengths across primates (16-18) – including in studies of dexterity (19). Additionally, we specifically demonstrate here that the metacarpal length for each finger is strongly and significantly associated with digit length (Figure S2, all parameters highly significant).

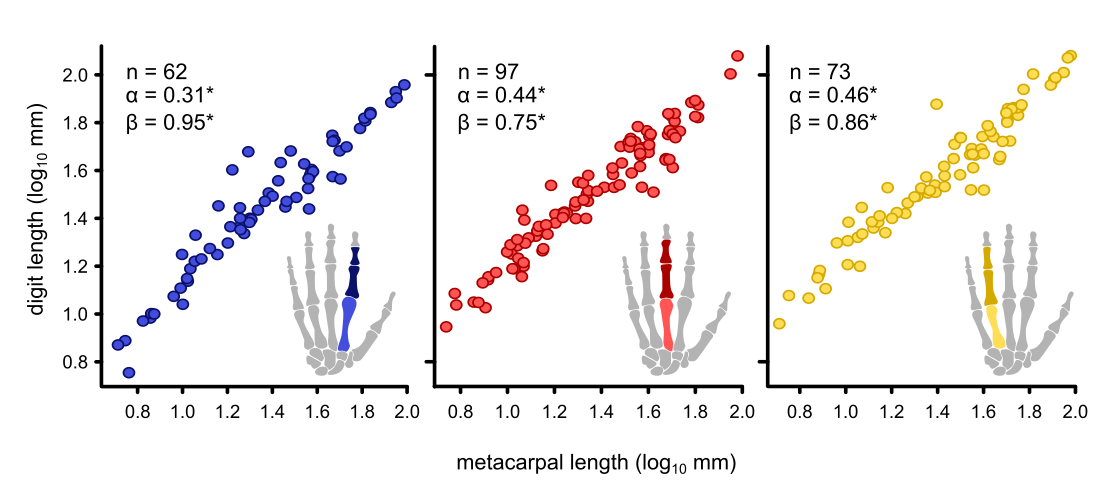

**Figure S2. The relationship between metacarpal length and digit length (measured as the sum of proximal phalanx and intermediate phalanx length) for digits 2-4.** Sample sizes, parameters and significance are indicated. *All parameters significant to p < 0.0001.

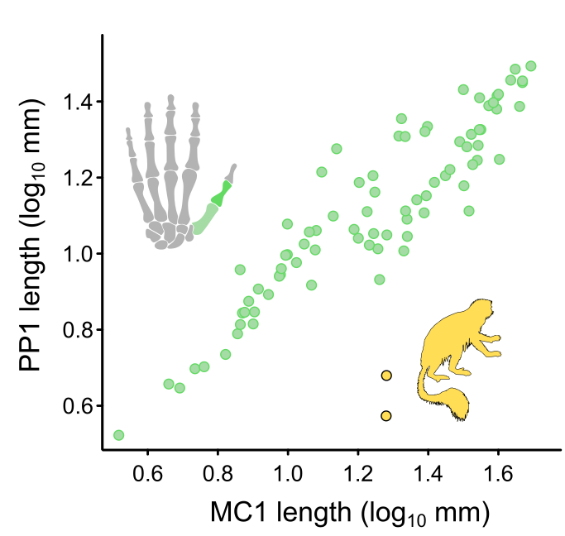
We calculated digit length as the sum of the lengths of the proximal phalanx and intermediate phalanx (excluding the metacarpal for comparisons). We exclude the distal phalanx owing to limited data availability. We then test the relationship between digit length and metacarpal length (both log_10_-transformed), using maximum-likelihood phylogenetic generalized least squares models implemented in caper (20), using a single representative phylogeny. We also find a significant relationship between the metacarpal and proximal phalanx of the thumb (Figure S3). Notable outliers are colobus monkeys, which (along with spider monkeys) are excluded from our analyses owing to their rudimentary thumbs (21).

**Figure S3. The relationship between the length of the metacarpal and the proximal phalanx in a sample of N = 175 primates with data for both bones.**

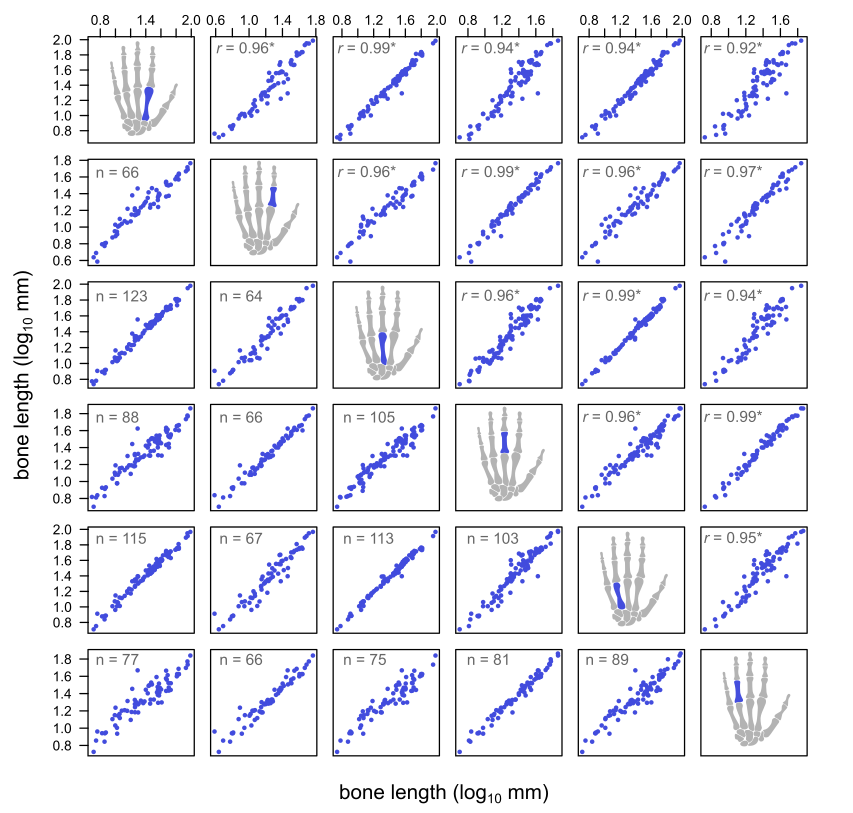
Additionally, there is a strong and significant correlation between all finger bones in our sample (digits 2-4, Figure S4).

**Figure S4. Pairwise correlation matrix between metacarpals and phalanges of the digits 2-4.** From top to bottom: MC2, PP2, MC3, PP3, MC4, PP4. Sample size for each comparison is shown in the lower half of the matrix and the correlation coefficient in the top half. All comparisons are highly significant, where * indicates p < 0.0001.

Finally, we find that our results are qualitatively identical when phalanges are used instead of metacarpals – or if we use the third or fourth digit to test the relationship between brain size and relative thumb length. The results of these tests are shown in Tables S1-S6.

**Table S1.** Parameter estimates for results using MC1 as the proxy for thumb length and MC2 as the proxy for finger length.

| **Model** | **N** | **R^2^** | **ϐ_[finger]_** | **N p_x_ < 0.05** | **Brain Region** | **ϐ_[brain]_** | **N p_x_ < 0.05** | **λ** |
| --- | --- | --- | --- | --- | --- | --- | --- | --- |
| finger-only | 94 | 0.82-0.84 | 0.87-0.89 | 100 | n/a | n/a | 100 | 0.82-0.89 |
| whole-brain | 94 | 0.85-0.86 | 0.69-0.72 | 100 | whole | 0.11-0.13 | 100 | 0.78-0.87 |
| finger-only | 88 | 0.85-0.87 | 0.89-0.91 | 100 | n/a | n/a | 100 | 0.84-0.91 |
| (excl. hominins) |  |  |  |  |  |  |  |  |
| whole-brain | 88 | 0.86-0.88 | 0.74-0.78 | 100 | whole | 0.08-0.11 | 100 | 0.84-0.91 |
| (excl. hominins) |  |  |  |  |  |  |  |  |

**Table S2.** Parameter estimates for results using MC1 as the proxy for thumb length and MC3 as the proxy for finger length.

| **Model** | **N** | **R^2^** | **ϐ_[finger]_** | **np_x_ < 0.05** | **Brain Region** | **ϐ_[brain]_** | **np_x_ < 0.05** | **λ** |
| --- | --- | --- | --- | --- | --- | --- | --- | --- |
| finger-only | 91 | 0.81-0.84 | 0.86-0.88 | 100 | n/a | n/a | 100 | 0.79-0.87 |
| whole-brain | 91 | 0.85-0.87 | 0.66-0.68 | 100 | whole | 0.13-0.14 | 100 | 0.72-0.84 |
| finger-only | 84 | 0.85-0.87 | 0.88-0.90 | 100 | n/a | n/a | 100 | 0.80-0.86 |
| (excl. hominins) |  |  |  |  |  |  |  |  |
| whole-brain | 84 | 0.86-0.88 | 0.71-0.74 | 100 | whole | 0.10-0.12 | 100 | 0.78-0.86 |
| (excl. hominins) |  |  |  |  |  |  |  |  |

**Table S3.** Parameter estimates for results using MC1 as the proxy for thumb length and MC4 as the proxy for finger length.

| **Model** | **N** | **R^2^** | **ϐ_[finger]_** | **np_x_ < 0.05** | **Brain Region** | **ϐ_[brain]_** | **np_x_ < 0.05** | **λ** |
| --- | --- | --- | --- | --- | --- | --- | --- | --- |
| finger-only | 106 | 0.77-0.81 | 0.83-0.86 | 100 | n/a | n/a | 100 | 0.88-0.97 |
| whole-brain | 106 | 0.82-0.86 | 0.63-0.68 | 100 | whole | 0.12-0.15 | 100 | 0.79-0.95 |
| finger-only | 99 | 0.87-0.89 | 0.87-0.91 | 100 | n/a | n/a | 100 | 0.85-0.95 |
| (excl. hominins) |  |  |  |  |  |  |  |  |
| whole-brain | 99 | 0.86-0.88 | 0.75-0.78 | 100 | whole | 0.08-0.10 | 100 | 0.86-0.95 |
| (excl. hominins) |  |  |  |  |  |  |  |  |

**Table S4.** Parameter estimates for results using PP1 as the proxy for thumb length and PP2 as the proxy for finger length.

| **Model** | **N** | **R^2^** | **ϐ_[finger]_** | **np_x_ < 0.05** | **Brain Region** | **ϐ_[brain]_** | **np_x_ < 0.05** | **λ** |
| --- | --- | --- | --- | --- | --- | --- | --- | --- |
| finger-only | 60 | 0.83-0.85 | 0.86-0.89 | 100 | n/a | n/a | 100 | 0.94-0.97 |
| whole-brain | 60 | 0.86 | 0.88 | 0.70-0.74 | whole | 0.11-0.13 | 100 | 0.91-0.95 |
| finger-only | 55 | 0.86-0.88 | 0.91-0.93 | 100 | n/a | n/a | 100 | 0.96-0.98 |
| (excl. hominins) |  |  |  |  |  |  |  |  |
| whole-brain | 55 | 0.87-0.89 | 0.75-0.79 | 100 | whole | 0.09-0.11 | 100 | 0.94-0.97 |
| (excl. hominins) |  |  |  |  |  |  |  |  |

**Table S5.** Parameter estimates for results using PP1 as the proxy for thumb length and PP3 as the proxy for finger length.

| **Model** | **N** | **R^2^** | **ϐ_[finger]_** | **np_x_ < 0.05** | **Brain Region** | **ϐ_[brain]_** | **np_x_ < 0.05** | **λ** |
| --- | --- | --- | --- | --- | --- | --- | --- | --- |
| finger-only | 72 | 0.80-0.84 | 0.84-0.86 | 100 | n/a | n/a | 100 | 0.78-0.89 |
| whole-brain | 72 | 0.84-0.86 | 0.66-0.70 | 100 | whole | 0.13-0.15 | 100 | 0.83-0.89 |
| finger-only | 66 | 0.86-0.87 | 0.87-0.89 | 100 | n/a | n/a | 100 | 0.71-0.80 |
| (excl. hominins) |  |  |  |  |  |  |  |  |
| whole-brain | 66 | 0.86-0.88 | 0.76-0.80 | 100 | whole | 0.07-0.09 | 98 | 0.76-0.85 |
| (excl. hominins) |  |  |  |  |  |  |  |  |

**Table S6.** Parameter estimates for results using PP1 as the proxy for thumb length and PP4 as the proxy for finger length.

| **Model** | **N** | **R^2^** | **ϐ_[finger]_** | **np_x_ < 0.05** | **Brain Region** | **ϐ_[brain]_** | **np_x_ < 0.05** | **λ** |
| --- | --- | --- | --- | --- | --- | --- | --- | --- |
| finger-only | 79 | 0.81-0.84 | 0.87-0.90 | 100 | n/a | n/a | 100 | 0.96-0.98 |
| whole-brain | 79 | 0.85-0.87 | 0.70-0.75 | 100 | whole | 0.11-0.14 | 100 | 0.94-0.97 |
| finger-only | 74 | 0.85-0.87 | 0.91-0.93 | 100 | n/a | n/a | 100 | 0.96-0.98 |
| (excl. hominins) |  |  |  |  |  |  |  |  |
| whole-brain | 74 | 0.86-0.88 | 0.77-0.81 | 100 | whole | 0.09-0.11 | 100 | 0.95-0.98 |
| (excl. hominins) |  |  |  |  |  |  |  |  |

### Peak workspace

Some authors argue that having a relatively long thumb does not necessarily lead to high manipulation ability and that we should instead quantify manipulation ability using kinematic models (12). In the kinematic model proposed by Feix et al. (12), a precision-grip *workspace* is calculated across a circular object of varying size scaled to hand size, resulting in a range of workspace values relative to a given object size. A ‘*manipulation workspace’* can be defined as the range of motion a small object can be freely moved between the thumb and index finger (12). One way of summarizing this is to look at the ‘peak workspace’, i.e. the object size at which a species (or specimen) has the highest workspace value.

We sought to demonstrate a significant association between brain size and peak workspace (accounting for object size) using a reduced sample of 41 primates with workspace data (12) – and accounting for the optimum object size which varies among species. To do this, we ran phylogenetic regression models in the same way as those we present in the main text. In these models, our response variable is the peak precision workspace and our predictor variables are brain size and object size. All variables are log_10_ transformed.

We find that brain size is a significant predictor of peak workspace (β_[brain]_ =0.09-0.11, p_x_<0.05 in 100% of topologies). The result is qualitatively identical when humans are excluded (β_[brain]_ =0.09-0.11, p_x_<0.05 in 100% of topologies). We also find a significant negative effect of object size on workspace (β_[object size]_ between -0.15 and -0.14, p_x_<0.05 in 100% of topologies).

Given these relationships, we therefore can conclude that peak workspace and relative thumb length are both reasonable proxies for overall manual dexterity across primates. Furthermore, there is a strong and significant association between the thumb length and peak workspace of each species. Including workspace as a covariate into our finger-only models demonstrates a highly significant association between the two variables (β_[workspace]_ =0.57-0.61, p_x_<0.05 in 100% of topologies) that is unaffected by the inclusion of humans or object size.

Whilst kinematic models are an elegant mechanical way to reconstruct and think about manipulative ability in individuals, they can be quite difficult to interpret or summarize into quantities that fit the expectations of evolutionary models. For example, modern humans have greater estimated manipulation potential than Neanderthals (12) – consistent with the idea that Neanderthals were more adapted for power gripping (22, 23) though see (24). However, the ‘optimal’ object size for humans is 19mm compared to the Neanderthal’s tiny 3mm. Biologically speaking, it is very unlikely that the hands of Neanderthals were under strong selection to specifically manipulate objects of 3mm in size.

### Brain Regions and Binocularity

We might expect that any improvements in fine-grained visuo-motor processes such as manual dexterity may have been driven by correlated increases in the regions of the brain responsible for both visual (e.g., the neocortex) and motor (e.g., the cerebellum) control.

The brain region data was taken from published literature (25-27) and consists of neocortex volume and cerebellum volume for a total of 49 species for which we have metacarpal data and whole brain sizes (see supplementary dataset S1). These are all extant taxa – we have no data for any hominins except modern humans.

In a subset of the whole-brain data for which we have data on the volume of both the neocortex and cerebellum (n = 49), we identified a significant positive relationship between both finger length (β_[finger]_ = 0.735, p_x_ < 0.05 in 100% of trees) and neocortex (β_[neocortex]_ = 0.178, p_x_ < 0.05 in 100% of trees, Figure 2b, Table 2). Surprisingly, there is no significant association found with the cerebellum in any model on any tree. Although correlated evolution of neocortex and cerebellum is a pronounced feature of primate brain evolution (28, 29), humans and other apes deviate from this pattern, exhibiting relative expansion of the cerebellum (25). However, there is no significant association between relative thumb length and cerebellum size regardless of the inclusion or exclusion of apes (and the sample size is too small to test in isolation, n = 6). The parameters of these models are presented in Table S7.

In our brain-region models, there is high phylogenetic signal (median λ = 0.955). Separating the effects of the neocortex and the cerebellum can be complicated owing to their strong correlation. Here, we include both regions in the same model here since together, the neocortex and cerebellum comprise a ‘unit’ responsible for the mediation of visuo-motor and sequential action control (30). However, we find qualitatively similar results when each of the regions are studied in isolation: without humans, only the neocortex shows any significant association (Table S8).

**Table S7.** Parameter estimates for results using brain regions as our response variable. The brain-regions models include both neocortex and cerebellum as well as finger length as a covariate. N= 49 for the model including *H. sapiens*.

| **Model** | **R^2^** | **ϐ_[finger]_** | **np_x_ < 0.05** | **Brain Region** | **ϐ_[brain]_** | **np_x_ < 0.05** | **λ** |
| --- | --- | --- | --- | --- | --- | --- | --- |
| brain-regions | 0.91-0.92 | 0.72-0.76 | 100 | neocortex | 0.16-0.20 | 100 | 0.92-0.97 |
|  |  |  |  | cerebellum | -0.04 | 0 |  |
| brain-regions | 0.91-0.93 | 0.79-0.82 | 100 | neocortex | 0.13-0.17 | 96 | 0.92-0.97 |
| (excl. H. sapiens) |  |  | 100 | cerebellum | -0.05 | 0 |  |

**Table S8.** Parameter estimates for results using only one of the brain regions as our response variable. These models include one of the two brain regions (neocortex and cerebellum) as well as finger length as a covariate. N= 49 for the model including *H. sapiens*.

| **Model** | **R^2^** | **ϐ_[finger]_** | **np_x_ < 0.05** | **ϐ_[brain]_** | **np_x_ < 0.05** | **λ** |
| --- | --- | --- | --- | --- | --- | --- |
| Neocortex | 0.91-0.92 | 0.70-0.73 | 100 | 0.12-0.14 | 100 | 0.94-0.98 |
| Neocortex (excl. H. sapiens) | 0.96 | 0.80-0.84 | 100 | 0.05-0.07 | 72 | 0.97-0.98 |
| Cerebellum | 0.90-0.92 | 0.73-0.77 | 100 | 0.10-0.12 | 100 | 0.91-0.97 |
| Cerebellum (excl. H. sapiens) | 0.96-0.97 | 0.85-0.89 | 100 | 0.03-0.05 | 0 | 0.97-0.98 |

Manipulation is visually guided. As brain size and binocularity has been linked to brain size evolution through the expansion of visual regions (31, 32), we additionally tested for an association with binocular field overlap and thumb length. Taking data from the literature on the degree of orbital convergence (33), we have 39 species that overlap with our thumb length and brain size data (see supplementary dataset S1). Note that this does not include any extinct taxa, nor is there data for *Homo sapiens.* We recover the expected association between brain size and binocularity (31): Both body size and binocularity are significant predictors of whole brain size in 100% of topologies ( β_body_ =0.55-0.56, β_binoc_ = 0.79-1.02). In terms of thumb length, the relationship with brain size is retained in this reduced sample, but there is no significant relationship with binocularity – either when studied in isolation or with brain size. In short, binocularity is associated with brain size, and so is thumb length – but they are not associated with one another.

We might expect that differences observed amongst primates might lead to non-significance in the cerebellum across primates. Strepsirrhines have relatively small neocortices in comparison to other primates (34). However, the results are qualitatively identical when we study the above relationships in strepsirrhines and haplorrhines separately.

All our results point to a singular conclusion: that neocortex size is implicated in thumb length evolution across primates, but not cerebellum size. However, further attention is required to untangle the specific effects of cortical regions on thumb length across primates.

### Dataset S1 (separate file).

The dataset associated with the study including lengths for the metacarpals, intermediate phalanges, proximal phalanges, brain mass, body mass, neocortex volume, and cerebellum volume. All references are included.

### Dataset S2 (separate file).

The sample of trees used in all analyses.

10.	Team RC. R: A Language and Environment for Statistical Computing. R Foundation for Statistical

Computing; 2024.
